# Remodeling-mediated changes in left ventricular mechanics under settings of chronic pressure overload and exercise

**DOI:** 10.1101/2025.08.09.669506

**Authors:** Francesco K. Yigamawano, Ricky Ruiz, Conner E. Johnson, Alexander Barnette, Lisa Freeburg, Kurt G. Barringhaus, Francis G. Spinale, Tarek Shazly

**Affiliations:** College of Engineering and Computing, Department of Biomedical Engineering, University of South Carolina, Columbia, South Carolina; Cardiovascular Translational Research Center, University of South Carolina School of Medicine, Columbia, South Carolina; Columbia VA Health Care System, Columbia, South Carolina; University of South Carolina School of Medicine Greenville, Greenville, South Carolina; School of Medicine, Department of Cell Biology and Anatomy, University of South Carolina, Columbia, South Carolina; Department of Cardiology, Prisma Health, Columbia, South Carolina

**Keywords:** left ventricle remodeling, myocardial mechanical properties, left ventricular chamber stiffness, speckle-tracking echocardiography, left ventricular pressure overload, exercise

## Abstract

Left ventricular (LV) remodeling, whether occurring as a part of somatic growth or as a chronic response to a sustained stimulus, is a primary factor underlying cardiac mechanical function. Although LV remodeling is a complex process that can be described at several levels (i.e. biochemical, cellular, tissue, organ, system), response variables that govern cardiac mechanics include changes in LV wall and chamber geometry, the mechanical properties of the LV myocardium, and functionally-deterministic LV structural mechanical properties such as LV chamber stiffness. In this study, we leverage two-dimensional speckle-tracking echocardiography (STE) to serially monitor key LV remodeling response variables in porcine models of LV pressure overload (LVPO), chronic exercise (CE), and the superposition of both settings (CE+LVPO), and compare changes to those occurring in age-matched referent control (RC) animals. Our findings show that over a 28-day study period, LVPO and CE both induce hypertrophy in comparison to RC, but passive LV myocardial stiffness increases with the former and decreases with the latter. As a net effect of geometrical and mechanical property changes, these settings induce divergent changes in LV chamber stiffness, namely an elevation with LVPO and reduction with CE. In the CE+LVPO cohort, exercise was found to attenuate the LVPO-induced increase in LV myocardial and LV chamber stiffnesses and insomuch supports its continued integration and refinement as a cardiac rehabilitation therapy. Data obtained across study cohorts were used to identify a phenomenological model of LV chamber stiffness, providing the first explicit relation of LV geometry and myocardial mechanical properties to a key LV structural property. Additional data processing was performed to develop a predictive mathematical model of late changes in LV chamber stiffness based on early remodeling response variables irrespective of stimulus, suggesting that STE can be extended to predict cardiac disease risk/progression in certain patient populations.

## 1. INTRODUCTION

Left ventricular (LV) remodeling is a multidimensional process in which cardiac mass, structure, and composition are progressively altered in response to sustained changes in hemodynamic load^1,2^. LV remodeling, which is a central part of normal physiological development, can be significantly influenced by persistent changes in hemodynamic load as manifest under settings of LV pressure overload (LVPO) and chronic exercise (CE)^3–5^. These settings differentially impact a wide range of LV remodeling response variables that cumulatively determine LV function and overall cardiac performance^6–8^.

Focusing on a subset of LV remodeling response variables with direct relevance to LV mechanics, analyses can be performed at two levels: (i) at the LV chamber-level, often captured through changes in LV chamber stiffness (or its inverse property, LV chamber compliance), LV mass, and LV chamber volumes and (ii) at the LV myocardial-level, commonly described by passive and active mechanical properties of the LV myocardium itself and the distribution of LV wall mass (i.e. regional LV wall thickness and inner radius). Changes in these key LV remodeling response variables are governed by mechanosensitive cellular and extracellular mechanisms involving cardiomyocyte hypertrophy, fibrosis, cytoskeletal remodeling, and extracellular matrix (ECM) production/reorganization^9–11^.

In both LVPO and CE settings, the LV myocardium is subjected to increased wall stress, which induces an adaptive hypertrophic response^12–14^. LV hypertrophy is *adaptive* in these settings as it tends to attenuate wall stress elevations in a manner first described by Y.C. Fung as the “principal of optimal mechanical operation”^15^. Although hypertrophy is an adaptive aspect of the LV remodeling response, additional changes – both at the LV chamber and LV myocardial levels – directionally differ in LVPO and CE settings and have direct implications for cardiac function.

Chronic LVPO, which can be induced by systemic hypertension or aortic stenosis, is associated with increased LV chamber stiffness and diastolic dysfunction. These adverse outcomes can be attributed to diffuse fibrosis with increased collagen synthesis, enhanced cross-linking, and a shift toward the stiffer (N2B) titin isoform^10,16–19^. Critically, the elevation in LV chamber stiffness with LVPO is a key process in the development of heart failure (HF) with preserved ejection fraction (HFpEF), which accounts for approximately 50% of all HF cases^20,21^. In contrast, in the setting of CE, LV remodeling results in reduced LV chamber stiffness and preserved/enhanced systolic function^22,23^. These changes are mediated by cardiomyocyte elongation, increased capillary density, and attenuation of profibrotic signaling cascades, including downregulation of transforming growth factor β (TGF-β) and modulation of matrix metalloproteinase (MMP) expression^24–26^.

LV chamber stiffness is thus a functionally deterministic property and promising clinical target for HFpEF therapies, including those with an exercise component. As a structural mechanical property, LV chamber stiffness is determined by LV myocardial stiffness as well as gross anatomical features such as the LV inner radius-to-wall thickness ratio and LV wall mass distribution. While the co-dependence of changes in LV chamber stiffness on the relative dynamics of geometrical and mechanical property changes is appreciated as a general concept, this structure-property-function relation has yet to be detailed in the context of normal cardiac development or under LVPO or CE settings. Moreover, the LV remodeling response under a superposition of these settings remains incompletely understood, although it is a key component underlying the success or failure of cardiac rehabilitation therapies based on prescribed exercise regiments^27,28^. Notably, animal studies examining related superimposed settings have yielded conflicting findings: in murine models, moderate CE following transverse aortic constriction (TAC) reduced fibrosis via suppression of the IL-6/JAK/STAT pathway, yet other studies reported worsened outcomes when CE was introduced in severe TAC settings^29–31^, underscoring the importance of timing, load severity, and myocardial reserve.

Motivated by gaps in fundamental knowledge on LV remodeling and the potential of CE as a widely accessible and affordable cardiac therapy, we performed a mechanically-focused investigation of LV remodeling in porcine models of (1) LVPO alone, (2) CE alone, and (3) CE superimposed on LVPO (CE+LVPO), as well as in (4) age-matched referent control (RC) animals. We hypothesized that with superimposed settings, CE would mitigate elevations in LV chamber stiffness induced by LVPO via a compensatory and dynamic reduction in LV myocardial stiffness. Obtained data provide novel insight into the magnitude and timing of LV remodeling-mediated regional changes in the LV myocardium in terms of LV geometry and LV myocardial mechanical properties, as well as global LV characteristics including LV ch8amber stiffness, LV mechanical heterogeneity, and LV mechanical anisotropy. In addition to these descriptive findings, our results enabled the interrelation of key geometrical and mechanical determinants of LV chamber stiffness in the context of a phenomenological model that is agnostic to the setting and thus representative of fundamental cardiac mechanics. Moreover, we identified an empirical equation to predict late changes in LV chamber stiffness based on the early LV remodeling trajectory, where again we found a common expression applicable across study cohorts. These findings yield new fundamental insights into LV remodeling and can inform tailored therapeutic approaches in cardiac rehabilitation programs.

## 2. METHODS

### 2.1 Study Cohorts

We investigated LV remodeling in porcine models of (1) LVPO, (2) CE, and (3) CE superimposed on LVPO (CE+LVPO), as well as in (4) age-matched referent control (RC) animals. All experimental procedures adhered to guidelines outlined by the National Institute of Health Guide for the Care and Use of Laboratory Animals and were approved by the Institutional Animal Care and Use Committee (IACUC) at the University of South Carolina. Protocols associated with each cohort have been extensively described in the referenced studies and are briefly summarized below. The current work builds upon previous findings by providing spatial and temporal resolution for LV remodeling response variables most germane to LV mechanics. A summary of obtained LV remodeling response variables is provided in Table 1.

**Table 1:** Abbreviations and Symbols.

|  |  |  |
| --- | --- | --- |
| LV | = | left ventricle |
| STE | = | speckle-tracking echocardiography |
| LVPO | = | left ventricle pressure overload |
| CE | = | chronic exercise |
| RC | = | referent control |
| ED | = | end-diastolic |
| LA | = | left atrial |
| EF | = | ejection fraction |
| HFpEF | = | heart failure with preserved ejection fraction |
| $K_c$ | = | left ventricle chamber stiffness |
| $\varepsilon_\theta; \varepsilon_r$ | = | circumferential myocardial strain; radial myocardial strain |
| $\sigma_\theta; \sigma_r$ | = | circumferential myocardial stress; radial myocardial stress |
| $K_{M,\theta}; K_{M,r}$ | = | circumferential myocardial stiffness; radial myocardial stiffness |
| $H_{Myo,\theta}; H_{Myo,r}$ | = | circumferential mechanical heterogeneity; radial mechanical heterogeneity |
| $A_{Myo}$ | = | mechanical anisotropy |

LVPO was induced in the pigs using a model previously described, where Yorkshire pigs (n=8, ∼25kg, castrated males) were anesthetized and underwent a left thoracotomy^8^. A 12-mm inflatable vascular cuff (Access Technologies, Skokie, IL) was secured around the supracoronary ascending aorta, with incremental inflation of the cuff each week over 4 weeks (initial inflation to 75 mmHg, increasing in 25 mmHg increments) to stimulate chronic LVPO. Post-surgical analgesia was provided using intramuscular buprenorphine (0.05 mg/kg) and fentanyl patches (50-100 mcg/hr). Previous studies demonstrated this model effectively induces LV hypertrophy, LV myocardial fibrosis, and elevated LV myocardial stiffness, with maintained ejection fraction (EF) that is consistent with clinical HFpEF phenotypes associated with LVPO^8,23,32^.

A second group of pigs (n=8, ∼25kg, castrated males) was subjected to a CE regimen as described previously, which included structured treadmill exercises (10 degrees incline, 2.5 mph, 10 minutes/day, 5 days/week over 4 weeks)^23^. Animals were acclimated for seven days prior to protocol to minimize stress during the procedure. The CE intervention aimed to evaluate the impact of physical activity on LV remodeling, previously demonstrating significant reductions in LV chamber stiffnesses and improved LV function by the study end-point^23,33–35^.

A third group of pigs (n=8, ∼25kg, castrated males) was subjected to the LVPO protocol described above while simultaneously participating in the treadmill exercise protocol (CE+LVPO). This setting superimposition was described in previous work and designed to assess whether regular physical activity mitigates the adverse LV remodeling effects of chronic pressure overload (Spinale FG, MD, PhD, submitted manuscript, 2025). Previous results showed that after 28 days of superimposed stimuli, LV remodeling resulted in muted elevation in LV myocardial stiffness, decreased LV myocardial fibrosis, and improved LV function compared to the LVPO-only group.

Age-, sex-, and weight-matched pigs (n=6) without LVPO or CE interventions were used as RCs for serial analysis across the 28-day study periods. This RC cohort facilitated comparative analyses and provided baseline normative data for LV structural and functional variables.

### 2.2 Serial Echocardiography

Two-dimensional speckle-tracking echocardiography (STE) was used to serially monitor LV remodeling response variables. Transthoracic echocardiography was performed using a GE Vivid E9 ultrasound system equipped with an M5S (1.5-4.6 MHz) transducer. Prior to imagining all animals were anesthetized using oral diazepam (200 mg) and intramuscular midazolam (0.5-0.6 mg/kg) to ensure minimal movement and stress. Serial echocardiographic images were obtained in short axis and long-axis views at baseline, weekly during the protocols, and post-protocol completion; doppler imaging was performed to enable calculation of LV end-diastolic pressure (EDP). Standard echocardiographic analyses and STE allowed for the assessment of LV wall/chamber geometry and LV myocardial deformation, respectively^36,37^.

STE was implemented on the short-axis echocardiographic view in prescribed LV myocardial regions, providing measures of regional LV myocardial strains in the circumferential (ε_θ_) and radial (ε_*r*_) direction based on relative changes in contained segmental lengths from end-systole (ES) to end-diastole (ED)^36,38^. The associated regional LV myocardial wall stresses at ED in the circumferential (σ_Θ_) and radial (σ_*r*_) directions were computed via a previously described methodology, which along with LV myocardial strains enabled calculation of regional LV myocardial stiffnesses in the circumferential (*K*_*M*,Θ_) and radial (*K*_*M*,*r*_) directions^8,37^. LV chamber stiffness (*K*_*C*_) was also computed in a serial fashion based on the EDP and the LV chamber volumetric strain using a previously described approach^23^.

### 2.3 LV Myocardial Heterogeneity and Anisotropy

The computed regional LV myocardial stiffnesses (*K*_*M*,Θ_ and *K*_*M*,*r*_) were used to derive indices of LV mechanical heterogeneity (*H*_*Myo*_) and anisotropy (*A*_*Myo*_). Heterogeneity – which reflects a spatial variation in material properties – was quantified in the circumferential (*H*_*Myo*,θ_) and radial (*H*_*Myo*,*r*_) directions based on the respective stiffness variance over all LV myocardial regions, respectively. Anisotropy – which reflects a directional difference in material properties – was quantified by A_Myo_ herein defined as the ratio of the LV myocardial stiffnesses (*K*_*M*,Θ_⁄*K*_*M*,*r*_) averaged over all LV myocardial regions.

### 2.4 Statistics

LV remodeling response variables measured weekly over the month-long protocols were analyzed using paired, two-tailed *t*-tests. Comparisons of response variables between age– and weight-matched animals across cohorts were performed using unpaired, two-tailed *t*-tests. Unless otherwise stated, all data are expressed as means ± standard error of the mean (SEM). Statistical analyses were conducted using MATLAB (MathWorks Inc., Natick, MA, USA) and Microsoft Excel (Microsoft Corporation, Redmond, WA, USA; 2018). p-values < 0.05 were considered statistically significant.

For the phenomenological model, model fit was assessed by the multiple R value (correlation coefficient) between the regression output and measured values. Similarly for the predictive model, performance was evaluated using the multiple R value; repeated measures (rm) analysis was applied to account for within-individual association and the paired structure of the dataset. The overall statistical significance of both models was evaluated using the significance F-value, which tests the null hypothesis that all regression coefficients are zero – i.e., that none of the independent variables explain the variation observed in the dependent variable.

## 3. RESULTS

### 3.1 End-diastolic LV Pressure, LV Myocardial Strain, and LV Myocardial Stress

Following our previously described approach for estimation of passive LV myocardial mechanical properties via STE-based analysis of diastolic filling^37^, we assume the following kinematics: at the onset-of-diastole, the LV is in the reference configuration and LV pressure, LV myocardial stress, and LV myocardial strain are all zero; at end-diastole, the LV myocardium is in a deformed configuration due to the end-diastolic pressure (EDP), with a deformation reflecting tensile circumferential strains/stresses and compressive radial strains/stresses.

While RC EDP remained relatively constant over the study period, LVPO caused a monotonic elevation of EDP culminating in a ∼31% increase by 28 days in comparison to the RC value. Conversely, CE induced a decrease in EDP with respect to RC at intermediate times (14-21 days post-protocol), and by day 28 EDP had reverted to the RC value. In the CE+LVPO cohort, EDP was elevated as with LVPO over 21 days post-protocol, but by day 28 the superposition of CE induced a reduction of ∼8% as compared to the LVPO alone value (Fig. 1A, Table 2).

**Figure 1:**
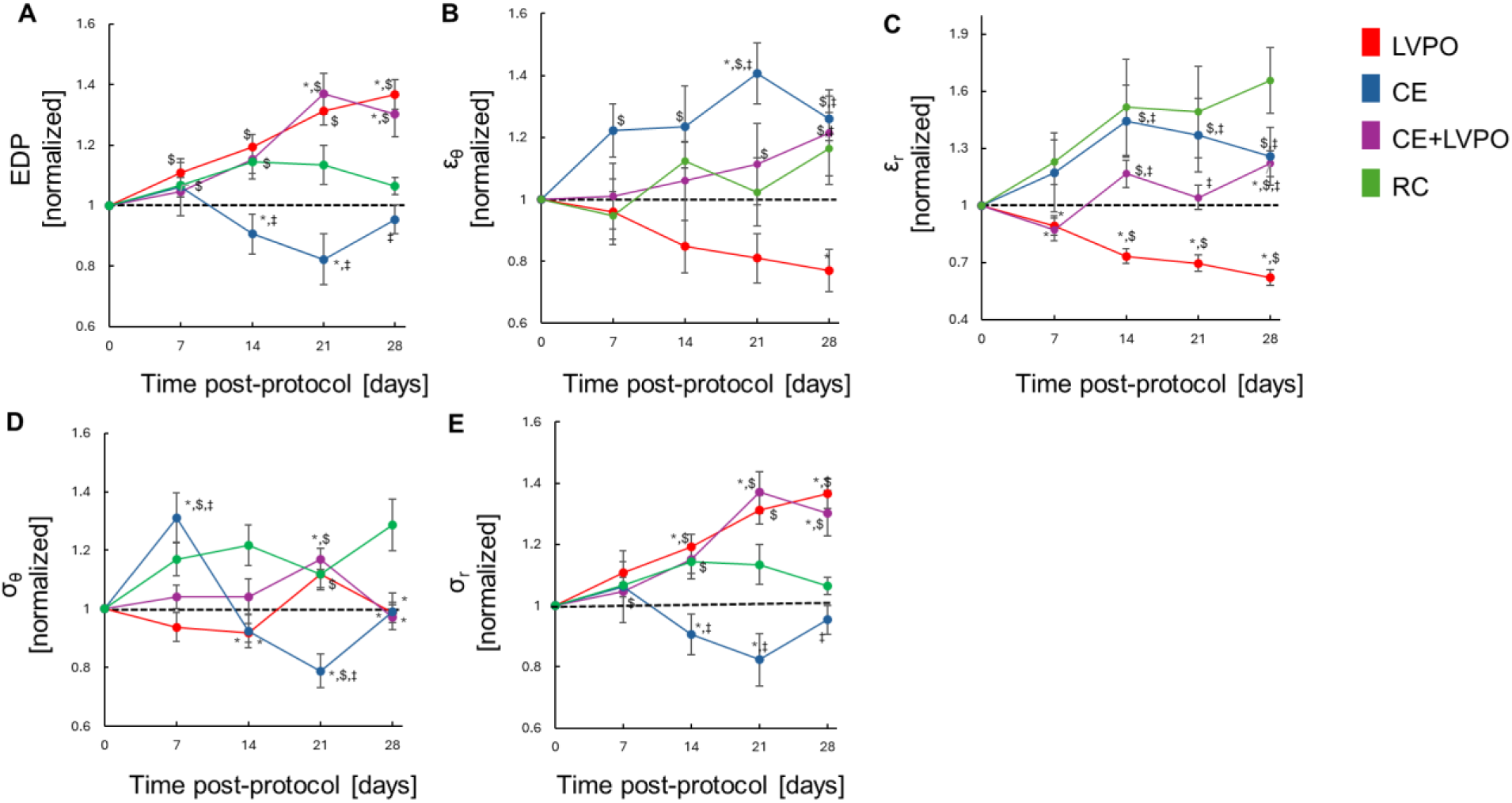
LV end-diastolic pressure and LV end-diastolic myocardial strains and stresses. **(A)** LV end-diastolic pressure (EDP), **(B)** circumferential LV myocardial strain (ε_θ_), **(C)** radial LV myocardial strain (ε_*r*_), **(D)** circumferential LV myocardial stress (σ_θ_), and **(E)** radial LV myocardial stress (σ_*r*_). All LV remodeling response variables are normalized with respect to individual baseline (day 0) values and presented as mean ± SEM. Study cohorts include LV pressure overload (LVPO), chronic exercise (CE), CE superimposed with LVPO (CE+LVPO), and referent control (RC). *indicates p < 0.05 vs. same day RC value; ^$^indicates p < 0.05 vs. same cohort baseline value; ^‡^indicates p < 0.05 vs same day LVPO value. n = 8 for LVPO and CE+LVPO cohorts; n = 6 for CE and RC cohorts.

**Table 2:** Initial and final (post-protocol day 28) values of LV remodeling response variables.

| **day 0**<br>**day 28** | **RC** | **LVPO** | **CE+LVPO** | **CE** |
| --- | --- | --- | --- | --- |
| **End-diastolic pressure [mmHg]** | 7.59 ± 0.34 | 8.05 ± 0.17 | 7.82 ± 0.32 | 8.64 ± 0.36 |
|  | 8.89 ± 0.45 | 10.97 ± 0.33<sup>*,§</sup> | 9.96 ± 0.20<sup>*,§</sup> | 8.17 ± 0.31 |
| **End-diastolic radial strain [-]** | -0.22 ± 0.01 | -0.28 ± 0.01 | -0.23 ± 0.01 | -0.22 ± 0.02 |
|  | -0.32 ± 0.01<sup>§</sup> | -0.18 ± 0.01<sup>*,§</sup> | -0.27 ± 0.01<sup>*,§</sup> | -0.26 ± 0.01<sup>*</sup> |
| **End-diastolic radial stress [kPa]** | -0.53 ± 0.02 | -0.54 ± 0.01 | -0.52 ± 0.02 | -0.58 ± 0.02 |
|  | -0.58 ± 0.03 | -0.73 ± 0.02<sup>*,§</sup> | -0.67 ± 0.03<sup>§</sup> | -0.54 ± 0.04 |
| **End-diastolic circumferential strain [-]** | 0.18 ± 0.02 | 0.19 ± 0.02 | 0.19 ± 0.02 | 0.16 ± 0.02 |
|  | 0.20 ± 0.02 | 0.15 ± 0.01<sup>*,§</sup> | 0.22 ± 0.02 | 0.20 ± 0.01<sup>§</sup> |
| **End-diastolic circumferential stress [kPa]** | 2.83 ± 0.17 | 3.38 ± 0.13 | 2.95 ± 0.11 | 3.70 ± 0.21 |
|  | 3.35 ± 0.11<sup>§</sup> | 3.27 ± 0.11 | 2.77 ± 0.09<sup>*</sup> | 3.38 ± 0.13 |
| **LV mass [g]** | 57.37 ± 5.87 | 56.79 ± 5.81 | 50.47 ± 2.27 | 48.97 ± 1.36 |
|  | 80.49 ± 7.12<sup>§</sup> | 114.04 ± 8.33<sup>*,§</sup> | 111.61 ± 2.43<sup>*,§</sup> | 98.14 ± 4.95<sup>§</sup> |
| **End-diastolic wall thickness [mm]** | 6.67 ± 0.61 | 5.81 ± 0.21 | 5.81 ± 0.37 | 6.17 ± 0.55 |
|  | 7.25 ± 0.50 | 10.43 ± 0.74<sup>*,§</sup> | 9.25 ± 0.57<sup>*,§</sup> | 8.16 ± 0.44 |
| **End-diastolic radius [mm]** | 15.67 ± 0.53 | 15.88 ± 0.34 | 15.91 ± 0.29 | 15.71 ± 0.66 |
|  | 18.33 ± 0.69<sup>§</sup> | 17.06 ± 0.48<sup>§</sup> | 17.72 ± 0.62<sup>§</sup> | 18.79 ± 0.51<sup>§</sup> |
| **End-diastolic volume [mL]** | 34.33 ± 3.33 | 31.38 ± 1.52 | 28.63 ± 2.01 | 30.67 ± 1.57 |
|  | 43.00 ± 2.70<sup>§</sup> | 44.82 ± 2.52<sup>§</sup> | 40.75 ± 2.53<sup>§</sup> | 53.50 ± 1.74<sup>§</sup> |
| **LV stroke volume [mL]** | 22.67 ± 2.49 | 21.69 ± 1.20 | 18.00 ± 1.33 | 19.00 ± 1.45 |
|  | 28.17 ± 1.79 | 31.19 ± 2.10<sup>§</sup> | 29.25 ± 3.12<sup>§</sup> | 37.33 ± 1.28<sup>*,§</sup> |
| **Ejection fraction [%]** | 65.41 ± 2.25 | 69.02 ± 1.21 | 63.08 ± 2.43 | 61.56 ± 2.31 |
|  | 65.64 ± 1.80 | 69.45 ± 1.93 | 71.08 ± 4.41 | 69.87 ± 1.74<sup>§</sup> |
| **Left atrial area [cm<sup>2</sup>]** | 4.75 ± 0.18 | 5.03 ± 0.32 | 5.39 ± 0.22 | 5.33 ± 0.24 |
|  | 7.30 ± 0.32<sup>§</sup> | 9.22 ± 0.43<sup>*,§</sup> | 7.80 ± 0.39<sup>§</sup> | 6.57 ± 0.25<sup>§</sup> |
| **LV chamber stiffness [kPa]** | 0.55 ± 0.06 | 0.48 ± 0.02 | 0.63 ± 0.07 | 0.74 ± 0.08 |
|  | 0.66 ± 0.06 | 0.70 ± 0.07<sup>§</sup> | 0.59 ± 0.12 | 0.47 ± 0.03<sup>*</sup> |
| **LV radial myocardial stiffness [kPa]** | 3.31 ± 0.44 | 2.29 ± 0.19 | 2.48 ± 0.12 | 3.46 ± 0.39 |
|  | 2.04 ± 0.08 | 6.18 ± 0.81<sup>*,§</sup> | 2.57 ± 0.08<sup>*</sup> | 2.38 ± 0.09 |
| **LV circumferential myocardial stiffness [kPa]** | 22.24 ± 3.02 | 23.36 ± 2.21 | 21.04 ± 2.21 | 31.22 ± 2.99 |
|  | 23.24 ± 2.99 | 42.44 ± 7.92<sup>*,§</sup> | 20.54 ± 2.05 | 19.20 ± 1.76<sup>§</sup> |

Values are presented as mean ± SEM. Sample sizes are n = 6 for RC and CE, n = 8 for LVPO and CE+LVPO. \*p < 0.05 vs. day 28 RC; §p < 0.05 vs. same cohort baseline.

The magnitudes of ε_θ_ and ε_*r*_monotonically decreased with LVPO, and by day 28 values were ∼30% and 60% lower than the respective RC values. In the CE cohorts LV myocardial strain magnitudes tended to increase over post-protocol times, but in comparison to RC there was an elevation in ε_θ_and reduction in ε_*r*_. In the CE+LVPO cohort, the trend in ε_θ_mirrored the RC cohort, while ε_*r*_magnitudes were comparatively reduced (Fig. 1B-C, Table 2).

Notable differences in LV myocardial stress trends relative to RC were observed in the CE cohort, whereby σ_Θ_ was comparatively elevated at early times (post-protocol day 7) and decreased at later times (post-protocol days 21-28). In terms of σ_*r*_, LVPO and CE+LVPO magnitudes were consistently higher than RC values over the study period, with ∼35% increase observed in these cohorts by post-protocol day 28 (Fig. 1D-E, Table 2).

### 3.2 LV Geometry and Function

LV hypertrophy – an increase in LV mass exceeding RC values – was expected and observed upon protocol completion in all experimental groups. By post-protocol day 28, normalized LV mass increases with LVPO, CE, and CE+LVPO were elevated relative to the RC cohort by ∼45%, 35%, and 39%, respectively (Fig. 2A, Table 2). Although the final extent of hypertrophy was similar in these groups, the rate of LV mass increase with CE was accelerated at later study times (post-protocol days 14-28), while in the LVPO and CE+LVPO cohorts the degree of hypertrophy was comparatively consistent over time.

**Figure 2:**
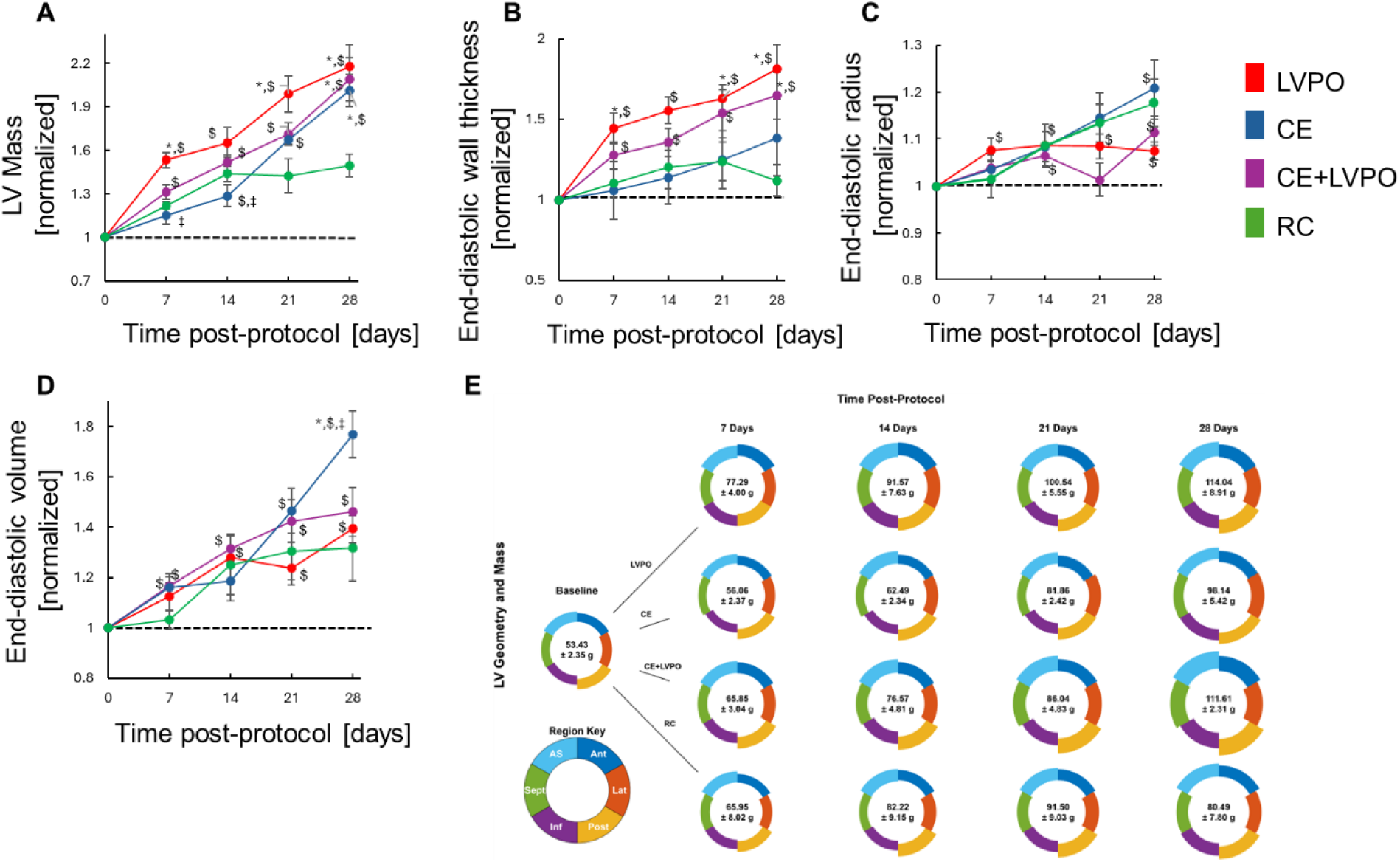
LV mass and geometry. **(A)** LV mass, **(B)** end-diastolic wall thickness, **(C)** end-diastolic radius, and **(D)** end-diastolic volume (EDV). All LV remodeling response variables are normalized with respect to individual baseline (day 0) values and presented as mean ± SEM. Study cohorts include LV pressure overload (LVPO), chronic exercise (CE), CE superimposed with LVPO (CE+LVPO), and referent control (RC). *indicates p < 0.05 vs. same day RC value; ^$^indicates p < 0.05 vs. same cohort baseline value; ^‡^indicates p < 0.05 vs same day LVPO value. **(E)** Schematic of regional LV geometry. Donut charts display the mean ED radius and ED wall thickness for six LV regions, namely anterior (Ant), lateral (Lat), posterior (Post), inferior (Inf), septal (Sept), and anterior septal (AS). The central value in each donut denotes total LV mass (mean ± SEM) at the indicated post-protocol time. Regional color keys are provided to aid interpretability. n = 8 for LVPO and CE+LVPO cohorts; n = 6 for CE and RC cohorts.

LV growth patterns underlying hypertrophy were further detailed via assessment of LV wall thickness and LV inner radius, both measured at ED, reported as mean values of six LV myocardial regions, and normalized with respect to cohort baseline values (Fig 2B-C). Consideration of trends in these geometrical variables suggests that with LVPO, early (post-protocol days 0-7) LV hypertrophy is due to increases in both LV wall thickness and inner radius (i.e. outward-hypertrophic), while analogous changes with CE manifest at later times in the LV remodeling process (over post-protocol days 21-28). In the CE+LVPO cohort, the final (post-protocol day 28) increases in LV wall thickness and LV inner radius fell between the values found for either setting alone, suggesting that exercise alters distribution of mass produced by LV remodeling. With respect to the progressive rise in EDV, the LVPO and CE+LVPO cohorts showed minor relative elevations (∼10%-15%) with respect to RC at post-protocol day 28, while in the CE cohort this relative elevation was more extreme (∼40%) and manifest by post-protocol day 21 (Fig. 2D, Table 2).

We further examined the geometrical outputs of LV remodeling via depiction of the regional growth throughout the LV myocardium (Fig. 2E). Obtained results qualitatively show that growth patterns are generally reflective of concentric hypertrophy in all cohorts, providing support for the previously described and herein utilized cylindrical cross-sectional model of the LV wall throughout the LV remodeling process.

In comparison to RC values, CE induced relative increases in LV stroke volume (SV) and ejection fraction (EF) by ∼30% and ∼15% respectively upon protocol completion if applied either alone or in combination with LVPO (Fig. 3A and B, Table 2). Both SV and EF were unaltered by LVPO alone, suggesting that the employed animal model recapitulates this defining feature of HFpEF development. The maximal value of left atrial (LA) area over the cardiac cycle was significantly elevated by LVPO and reduced by CE, while changes in the CE+LVPO cohort tracked closely with RC (Fig. 3C, Table 2).

**Figure 3:**
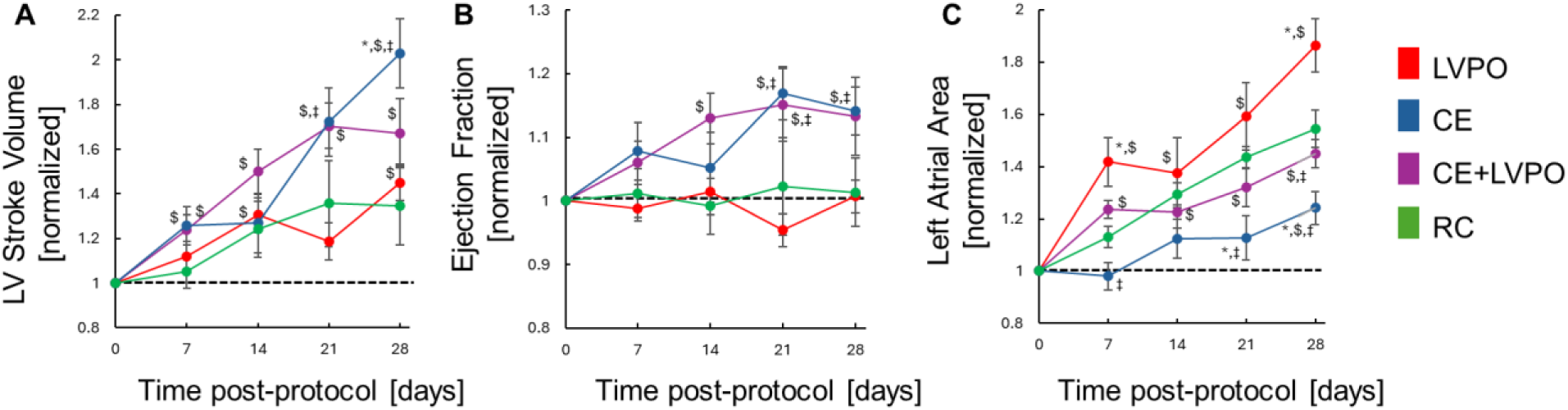
LV function. **(A)** LV stroke volume, **(B)** ejection fraction, and **(C)** left atrial area. All LV remodeling response variables are normalized with respect to individual baseline (day 0) values and presented as mean ± SEM. Study cohorts include LV pressure overload (LVPO), chronic exercise (CE), CE superimposed with LVPO (CE+LVPO), and referent control (RC). *indicates p < 0.05 vs. same day RC value; ^$^indicates p < 0.05 vs. same cohort baseline value; ^‡^indicates p < 0.05 vs same day LVPO value. n = 8 for LVPO and CE+LVPO cohorts; n = 6 for CE and RC cohorts.

### 3.3 LV Myocardial Stiffness and LV Chamber Stiffness

LV remodeling-mediated changes in *K*_*M*,Θ_ and *K*_*M*,*r*_ diverged in LVPO and CE settings, with values progressively increasing in the former and decreasing with the latter (Fig. 4A and B, Table 2). At LVPO post-protocol day 28, *K*_*M*,Θ_ and *K*_*M*,*r*_ were elevated with respect to baseline values by ∼80% and 140%, respectively, while with CE there were analogous reductions of ∼35% and 25%. In the CE+LVPO cohort, these opposing trends were largely offsetting, resulting in post-protocol day 28 values of both *K*_*M*,Θ_ and *K*_*M*,*r*_ that were statistically indistinguishable from baseline values. Trends in *K*_*C*_ were similar across cohorts at protocol completion, with LVPO inducing a post-protocol day 28 increase of ∼50% with respect to baseline, and CE an analogous decrease of ∼35% (Fig. 4C, Table 2). As with LV myocardial stiffness, the superposition of stimuli offset these opposing trends, resulting in a retention of baseline *K*_*C*_ throughout the CE+LVPO study period (Fig. 4C, Table 2).

**Figure 4:**
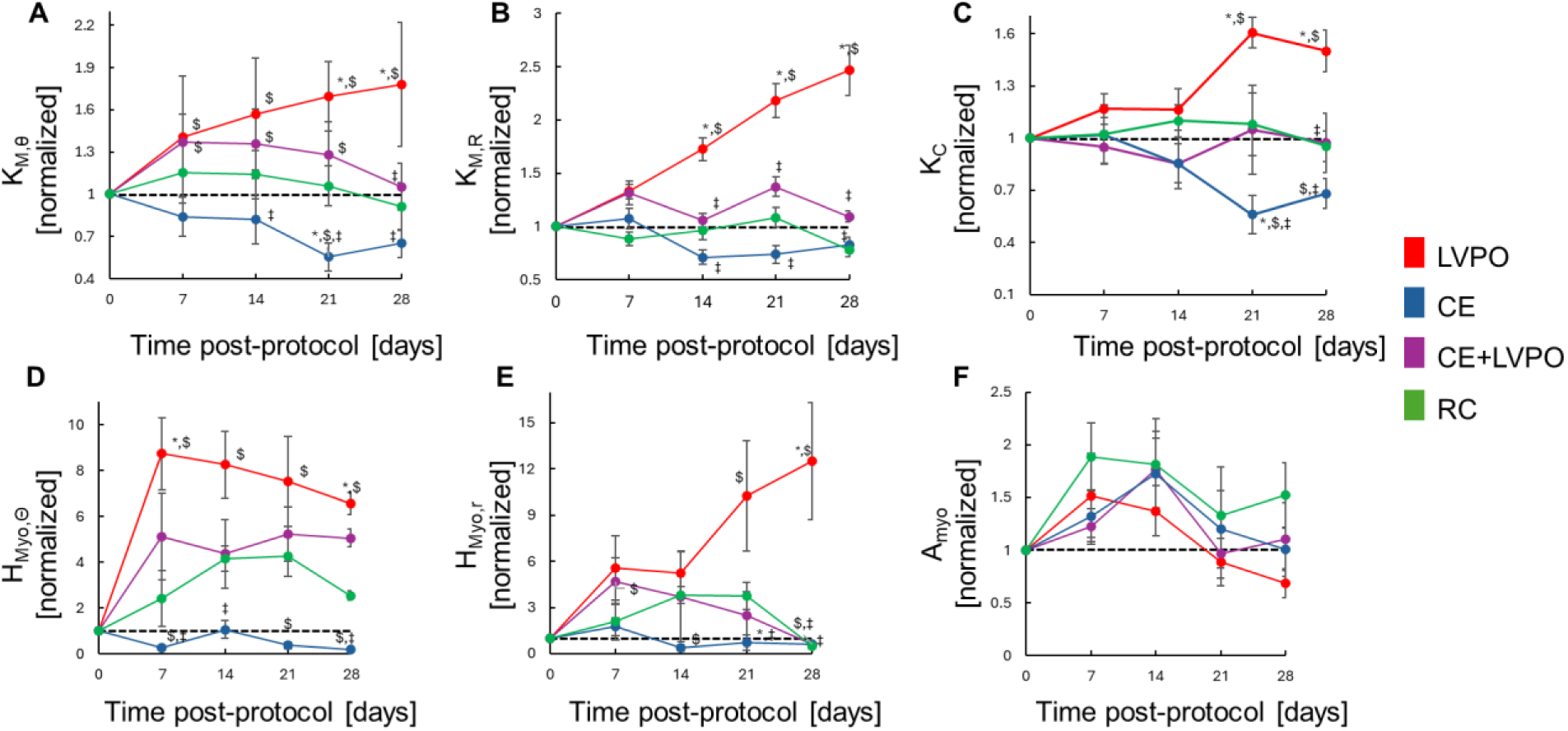
LV structural and mechanical properties. **(A)** LV myocardial circumferential stiffness (*K*_*M*,θ_), **(B)** LV myocardial radial stiffness (*K*_*M*,*r*_), **(C)** LV chamber stiffness (*K*_*C*_), LV mechanical heterogeneity with respect to **(D)** circumferential (*H*_*Myo*,θ_) and **(E)** radial LV myocardial (*H*_*Myo*,*r*_) stiffnesses, and **(F)** LV mechanical anisotropy (*A*_*Myo*_). All LV remodeling response variables are normalized with respect to individual baseline (day 0) values and presented as mean ± SEM. Study cohorts include LV pressure overload (LVPO), chronic exercise (CE), CE superimposed with LVPO (CE+LVPO), and referent control (RC). *indicates p < 0.05 vs. same day RC value; ^$^indicates p < 0.05 vs. same cohort baseline value; ^‡^indicates p < 0.05 vs same day LVPO value. n = 8 for LVPO and CE+LVPO cohorts; n = 6 for CE and RC cohorts.

### 3.4 LV Heterogeneity and Anisotropy

LV remodeling in the LVPO setting resulted in a significant increase in LV myocardial heterogeneity, wherein post-protocol day 28 *H*_*Myo*,θ_ and *H*_*Myo*,*r*_ values were elevated by ∼7-fold and 12-fold, respectively, relative to baseline values (Fig. 4D and E, Table 2). Conversely, CE promoted the retention of baseline LV myocardial heterogeneity, with no significant changes in either *H*_*Myo*,θ_ or *H*_*Myo*,*r*_ throughout the study period. In the CE+LVPO cohort, trends in both *H*_*Myo*,θ_ and *H*_*Myo*,*r*_ exhibited no statistical difference from RC and again suggested an offsetting effect with stimuli superposition.

*A*_*Myo*_ exhibits a nonmonotonic change in all four study groups, generally characterized by an early (post-protocol days 0-14) increase followed by a late (post-protocol days 14-28) decrease towards the baseline values (Fig. 4F, Table 2). Although observed differences among cohorts do not rise to the level of statistical significance, comparison among trends suggest that LVPO reduces the degree of anisotropy relative to RC, culminating with a normalized *A*_*Myo*_ < 1 reflecting an enhanced intraregional rate of increase in *K*_*M*,*r*_ in comparison to *K*_*M*,Θ_.

### 3.5 Phenomenological and Predictive Models of LV Chamber Stiffness

Obtained data were used to identify both a phenomenological and predictive model of *K*_*C*_. Both models are empirically-determined linear equations that are applicable across study cohorts and insomuch reflect a fundamental relation governing LV mechanics. The objective of the phenomenological model is to explicitly relate *K*_*C*_ to a subset of LV remodeling response variables that quantify LV geometry and myocardial mechanical properties, providing a structure-function-property relation that interrelates key variables in LV remodeling. The predictive model aims to utilize data obtained early in the LV remodeling process (post-protocol days 0-14) to predict late (post-protocol day 28) values of *K*_*C*_, thereby providing a quantitative basis for evaluating HFpEF trajectory and/or the efficacy of CE-based cardiac therapy.

The resultant phenomenological model (Eq. 1) contains two dimensionless constants (a, b; Table 3) that modulate a linear dependence of *K*_*C*_ on end-diastolic wall thickness (t), ESV, and LV myocardial stiffnesses (*K*_*M*,Θ_ and *K*_*M*,*r*_) and an inverse dependence on mean end-diastolic inner radius (*r*_*i*_) and EDV. Mean values across LV myocardial regions were used for all model variables with regional variance (t, *r*_*i*_, *K*_*M*,Θ_ and *K*_*M*,*r*_). Modeled values showed reasonable agreement (R=0.855, F(2, 139) < 0.001) with direct measures of *K*_*C*_ (Fig. 5A) across all study cohorts/post-protocol times.

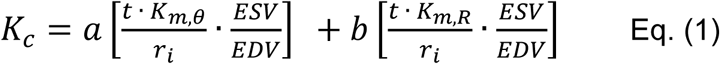

**Figure 5:**
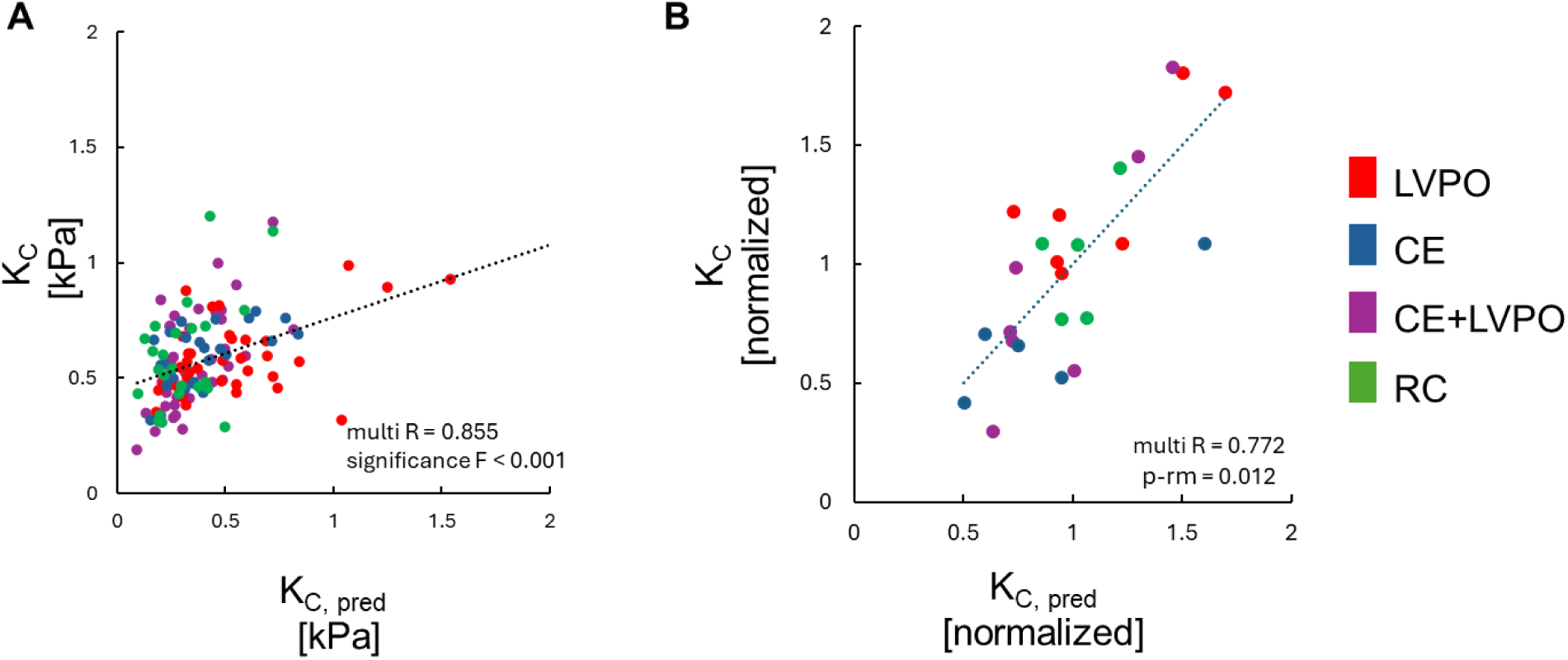
Phenomenological and predictive models of LV chamber stiffness. **(A)** A stepwise multiple-regression model (Eq. 1) was used to identify a phenomenological relation between LV chamber stiffness (*K*_*C*_) and concurrent measures of LV mechanical properties and LV geometry (Eq. 1), with utilization of data from all study cohorts. **(B)** A stepwise multi-linear regression model (Eq. 2.) was used to predict the post-protocol day 28 *K*_*C*_ based on early-stage (post-protocol days 0-14) changes of LV remodeling response variables. For the predictive model, *K*_*C*_ is normalized with respect to individual baseline (day 0) values.

**Table 3:** Model fitting parameters.

|  | Parameters |  |  |  |
| --- | --- | --- | --- | --- |
|  | a | b | c | d |
| Phenomenological model (Eq. 1) | 0.0963 | 0.0587 | - | - |
| Predictive model (Eq. 2) | 1.06 | -0.0293 | -0.184 | 1.05 |

The resultant predictive model (Eq. 2) contains four dimensionless constants (a, b, c, d; Table 3) that reflect a proportionality between the normalized value of *K*_*C*_ at post-protocol day 28 and the percentage change from baseline to post-protocol day 14(%Δ0−14) of a subset of LV remodeling response variables, namely 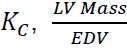, and ESV. Predicted values showed reasonable agreement (R=0.772, p-rm= 0.012, F(3,22) < 0.001) with direct measures of *K*_*C*_ (Fig. 5B) across all study cohorts.

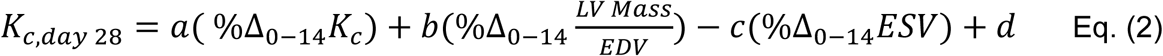

## 4. DISCUSSION

### 4.1 Summary of the LV Remodeling Response in Diverse Settings

Our findings delineate LV remodeling trajectories induced by LVPO and CE in isolation or in combination, emphasizing the divergent trends in mechanical and geometric response variables. To frame our understanding of LV remodeling under the diverse settings, our study includes a detailed assessment of progressive changes of key response variables in RC animals. Our RC data supports the notion of preferred/homeostatic LV myocardial stress levels and their retention during development (Fig. 1D and E, Table 2), as σ_Θ_ and σ_*r*_ magnitudes at ED were ∼3 kPa and 0.5 kPa, respectively, throughout the study period^39,40^. Interestingly, in all other considered settings (LVPO, CE, CE+LVPO), σ_Θ_ was similarly retained, but σ_*r*_ was progressively elevated in the LVPO setting with or without CE (Fig. 1D and E, Table 2). This implies that irrespective of the setting, LV remodeling maintains an adaptive character in terms of the retaining the homeostatic stress state of mechanosensitive cardiac cells in their primary direction of the alignment – the circumferential direction^41,42^.

Consistent with prior literature, LVPO led to an elevation in EDP, progressive hypertrophy mainly due to an increase in wall thickness, and a substantial rise in passive LV myocardial stiffness and LV chamber stiffness^8,43^ (Fig. 4A-C, Table 2). These changes were simultaneous with a stark increase in LV myocardial heterogeneity and a tendency to mute LV anisotropy (Fig 4D-F, Table 2), further underscoring the complexity in changes underlying the deleterious nature of LVPO-induced remodeling. Conversely, CE induced hypertrophy coupled with other favorable adaptations, including a reduction in both LV myocardial and chamber stiffnesses (Fig. 4A-C, Table 2). Notably, CE preserved or enhanced EF and SV, reduced EDP, and preserved RC trends in LV myocardial heterogeneity and anisotropy throughout the LV remodeling process (Fig 4D-F, Table 2).

### 4.2 Support for CE in Cardiac Therapy

The superimposed CE+LVPO setting produced an intermediate LV remodeling response, as response variables typically fell between values seen with either setting in isolation. Critically, LVPO-driven elevations in LV myocardial stiffness and LV chamber stiffness were effectively attenuated (Fig. 4A-C, Table 2), and except for LV mass, the time-course and magnitude of LV remodeling response variables often reflected RC trends. These unique data provide mechanistic insight into the therapeutic benefits of CE in mitigating adverse structural and mechanical remodeling induced by chronic LVPO^44^, with quantification of changes in LV myocardial stiffness and LV chamber stiffness highlighting their effect on fundamental determinants of diastolic function^45,46^. These findings are particularly relevant in the context of HFpEF, wherein increased *K*_*C*_is a hallmark pathological feature. The CE-induced reversal of key maladaptive mechanical signatures highlights its potential as a cornerstone in cardiac rehabilitation programs for HFpEF patients. Furthermore, the preservation of EF and SV in this cohort suggests maintenance of systolic function, an essential clinical outcome. The observed ability of CE to blunt LVPO-induced alterations in LV myocardial heterogeneity and anisotropy (Fig. 4 D-F) further supports its role in promoting uniform myocardial adaptation and balanced mechanical function. Given the relationship between regional mechanical heterogeneity and arrhythmogenic risk, CE’s effect in preserving electromechanical synchrony may also carry implications for arrhythmic burden reduction^47^.

### 4.3 Utility of STE to Evaluate and Refine Therapeutic Protocols

This study demonstrates the utility of STE as a powerful modality for monitoring LV remodeling and evaluating therapeutic interventions. STE enabled high-resolution, regionally-specific quantification of LV myocardial strain and stress, allowing for robust derivation of both LV myocardial stiffness and LV chamber stiffness over time. Such temporal and spatial resolution is essential in capturing the dynamic interplay between LV myocardial mechanics and geometry during LV remodeling.

Notably, we were able to identify a linear phenomenological model relating *K*_*C*_ to key structural and mechanical variables across cohorts. This model not only reflects fundamental mechanical principles governing chamber stiffness but may serve as a translational tool for non-invasive estimation of *K*_*C*_ in clinical populations. Its simplicity and generalizability are strengths that support its use in real-time patient monitoring. Equally significant is the predictive model we developed, which utilized early changes in remodeling variables (by day 14) to forecast terminal values of *K*_*C*_ (by day 28). This capability suggests that STE-derived variables may be valuable in prognostication and early stratification of patients at risk for maladaptive remodeling, providing a window for early therapeutic intervention and protocol optimization. In this way, STE becomes more than a diagnostic tool—it evolves into a platform for dynamic therapy guidance.

### 4.4 Study Limitations

Despite the valuable insights gained, several limitations should be acknowledged. First, while the porcine model provides a close approximation of human cardiac physiology, interspecies differences remain and may affect the translation of our findings. Second, although the study duration captured the early to intermediate phases of LV remodeling, longer-term studies are necessary to assess whether the beneficial effects of CE persist or evolve over time. Third, while we focused on passive myocardial mechanical properties, active contractile parameters and molecular markers (e.g., fibrosis, titin isoforms, MMP/TIMP expression) were not herein evaluated and may provide further insight into the observed mechanical adaptations. Additionally, the moderate intensity and short duration of the CE protocol may limit extrapolation to other exercise regimens or clinical populations with advanced HFpEF. Future studies should evaluate dose-response relationships of CE, the role of timing relative to disease progression, and the reversibility of established pathological remodeling.

## 5. CONCLUSION

This study presents a comprehensive, mechanics-focused characterization LV remodeling in response to LVPO, CE, and their superimposition, using porcine models and high-resolution STE. Our results demonstrate that LVPO and CE independently induce hypertrophy but lead to divergent structural and functional outcomes over time— namely, progressive increases in LV myocardial and LV chamber stiffness with LVPO and analogous decreases with CE. Importantly, CE effectively attenuated the maladaptive aspects of mechanical changes associated with LVPO when the stimuli were combined, restoring LV myocardial stiffness and LV chamber stiffness to near baseline values. Our findings reinforce the value of CE as a therapeutic strategy in conditions characterized by elevated LV chamber stiffness, such as HFpEF. Moreover, the study introduces a novel phenomenological model that relates key structural and mechanical variables to LV chamber stiffness, as well as a predictive model that forecasts late remodeling outcomes based on early indicators. Together, these models offer a translational framework for non-invasive monitoring and early risk stratification in cardiac disease. Taken together, our findings deepen mechanistic understanding of LV remodeling, validate the use of STE as a dynamic and predictive tool in both experimental and clinical settings, and underscore the promise of exercise-based interventions in mitigating pathological cardiac remodeling.

## ACKNOWLEDGEMENTS

None

